# Sex differences in predictors and regional patterns of brain-age-gap estimates

**DOI:** 10.1101/2022.01.19.476964

**Authors:** N Sanford, R Ge, M Antoniades, A Modabbernia, SS Haas, HC Whalley, L Galea, SG Popescu, JH Cole, S Frangou

## Abstract

**Background:** The brain-age-gap estimate (brainAGE) quantifies the difference between chronological age and age predicted by applying machine-learning models to neuroimaging data, and is considered a biomarker of brain health. Understanding sex-differences in brainAGE is a significant step toward precision medicine.

**Methods:** Global and local brainAGE (G-brainAGE and L-brainAGE, respectively) were computed by applying machine learning algorithms to brain structural magnetic resonance imaging data from 1113 healthy young adults (54.45% females; age range: 22-37 years) participating in the Human Connectome Project. Sex-differences were determined in G-brainAGE and L-brainAGE. Random forest regression was used to determine sex-specific associations between G-brainAGE and non-imaging measures pertaining to sociodemographic characteristics and mental, physical, and cognitive functions.

**Results:** L-brainAGE showed sex-specific differences in brain ageing. In females, compared to males, L-brainAGE was higher in the cerebellum and brainstem and lower in the prefrontal cortex and insula. Although sex-differences in G-brainAGE were minimal, associations between G-brainAGE and non-imaging measures differed between sexes with the exception for poor sleep quality, which was common to both. The most important predictor of higher G-brainAGE was non-white race in males and systolic blood pressure in females.

**Conclusions:** The results demonstrate the value of applying sex-specific analyses and machine learning methods to advance our understanding of sex-related differences in factors that influence the rate of brain ageing and provide a foundation for targeted interventions.

## Introduction

Neuroimaging studies have been instrumental in identifying sex-differences in brain structure across the lifespan (Ge et al., 2021; Jahanshad & Thompson, 2017; Kaczkurkin et al., 2019; Ruigrok et al., 2014). Brain structure shows profound age-related changes throughout the lifespan (Dima et al., 2021; Frangou et al., 2021; Lebel & Beaulieu, 2011; Storsve et al., 2014; Tamnes et al., 2013; Wierenga et al., 2020), which are also modified by sex. Females show somewhat accelerated brain maturation during adolescence, suggesting a link with pubertal onset (Ball et al., 2021; Brouwer et al., 2021). In middle and late adulthood, sex differences in age-related brain changes appear less pronounced (Bittner et al., 2021) but female brains may retain more “youthful” transcriptomic and metabolic features than male brains (Beheshti et al., 2021; Berchtold et al., 2008; Goyal et al., 2019; Goyal et al., 2017; Skene et al., 2017). In females, there are fewer age-related changes in in aerobic glycolysis (Goyal et al., 2019; Goyal et al., 2017) and in the expression of genes related to energy production and protein synthesis (Berchtold et al., 2008; Skene et al., 2017). These transcriptomic and metabolic findings show regional differences, with females retaining more youthful features primarily in prefrontal areas (Beheshti et al., 2021; Goyal et al., 2017). Whether this pattern is also reflected in macro-structural brain morphometry is currently unknown.

Besides sex, numerous factors are known to influence the rate of age-related brain changes. Those thought to accelerate brain ageing notably include smoking (Karama et al., 2015), alcohol use (Topiwala et al., 2017), obesity (Gurholt et al., 2021), hypertension (Gurholt et al., 2021), psychopathology (Wertz et al., 2021), poor inter-personal function (Hatton et al., 2018) and lower socioeconomic status (Chan et al., 2018). Conversely, the rate of brain ageing may be attenuated in individuals with higher cognitive function and educational attainment (Elliott et al., 2019) and better physical fitness (Steffener et al., 2016). Associations between accelerated brain ageing and indicators of age-dependent decline can be detected as early as the 3^rd^ and 4^th^ decade of life, even in generally healthy individuals (Belsky et al., 2015; Elliott et al., 2019). This evidence underscores the importance of focusing on young adulthood while testing for sex differences in the relative importance of risk, as attenuating factors in this age-group may contribute towards targeted interventions for preventing deterioration in brain health later in life (Belsky et al., 2015; Elliott et al., 2019).

Based on these considerations, the current study aims to advance knowledge on sex differences in the neuroanatomical pattern and in the predictors of brain ageing in young adulthood. To achieve this, we availed of machine learning methods to yield estimates of the biological age of the brain (brain-age) based on neuroimaging features. The gap between brain-age and chronological age, referred to here as the brain-age-gap estimate (brainAGE), enables inferences about the apparent acceleration or delay in age-related biological processes. We used structural magnetic resonance imaging (MRI) data obtained from young adults (age range 22-37 years) participating in the Human Connectome Project (HCP; https://www.humanconnectome.org/) to compute global (G-brainAGE) and local brainAGE (L-brainAGE). G-brainAGE is a global index of age-related changes across the brain (Cole & Franke, 2017; Cole et al., 2017; Franke et al., 2010) while L-brainAGE informs about regional age-related brain changes (Popescu et al., 2021). We hypothesized that sex differences in L-brainAGE might follow the same pattern observed in transcriptomic and metabolic data (Beheshti et al., 2021; Berchtold et al., 2008; Goyal et al., 2019; Goyal et al., 2017; Skene et al., 2017), with females having more youthful brains particularly in prefrontal regions. The HCP also includes non-imaging variables (NIMs) pertaining to demographic characteristics, cognition, mental health, interpersonal relationships, physical fitness, and lifestyle characteristics that enable testing for sex differences in factors that may accelerate or protect against age-related brain changes.

## Methods

### Participants

We used data from the S1200 public release of the Human Connectome Project (HCP; https://www.humanconnectome.org/study/hcp-young-adult/document/1200-subjects-data-release) which comprises 1113 healthy young adults (606 females) with a mean age of 28.80 years (range 22–37 years).

### Neuroimaging

Whole-brain T_1_-weighted magnetic resonance imaging (MRI) scans were acquired in the HCP participants on a Siemens Skyra 3T scanner (Erlanger, Germany) (details in supplemental material). Images were downloaded from the HCP repository and processed locally.

#### Local brainAGE computation

The process for generating local brainAGE estimates followed the procedures developed by Popescu and colleagues (2021), which are described in the supplemental material. Briefly, the T_1_-weighted images of the HCP participants were normalized using affine followed by non-linear registration, corrected for bias field inhomogeneities, and segmented into gray and white matter and cerebrospinal fluid components. The Diffeomorphic Anatomic Registration Through Exponentiated Lie algebra algorithm (DARTEL; Ashburner, 2007) was applied to normalize the segmented scans into a standard MNI space (MNI-152 space). The gray and white matter outputs were used as input to a pre-trained, convolutional neural network (U-Net) to yield voxel-wise estimates of L-brainAGE using parameters provided by Popescu and colleagues (2021) (https://github.com/SebastianPopescu/U-NET-for-LocalBrainAge-prediction). The model was trained and tested in a sample of T_1_-weighted brain scans from 4,155 healthy individuals aged 18-90 years. Importantly, the HCP dataset was not used in the development of the L-brainAGE model. Performance accuracy was ascertained using the voxel-level Mean Absolute Error (MAE), which quantifies the absolute difference between the neuroimaging-predicted age and the chronological age. The MAE was not adjusted for chronological age

#### Global brainAGE computation

Downloaded T_1_-weighted images for HCP participants were processed using standard pipelines implemented in SPM12 (https://www.fil.ion.ucl.ac.uk/spm/software/spm12/) and the computational anatomy toolbox (CAT12 version r1155; Gaser & Dahnke, 2016) (http://www.neuro.uni-jena.de/cat/). The output was used as the input features in a linear support vector regression with 10-fold cross-validation (Schölkopf & Smola, 2002) implemented in the freely available machine-learning software NeuroMiner (https://github.com/neurominer-git/NeuroMiner-1), which has been widely used for age prediction from neuroimaging data (Besteher et al., 2019; Koutsouleris et al., 2014; Löwe, et al., 2016) (details in supplemental material). The age-prediction models were conducted separately for male and female HCP participants. The MAE was used to ascertain model accuracy and was not adjusted for chronological age. G-brainAGE in each HCP participant was computed by subtracting their chronological age from the neuroimaging-predicted age. To account for residual associations with age, G-brainAGE was corrected separately in males and females by regressing out chronological age as per Le et al. (2018); unless otherwise specified, the corrected G-brainAGE values were used in all subsequent analyses.

For both L-brainAGE and G-brainAGE, positive values indicate higher brain-age relative to chronological age, while the opposite is true for negative values.

### Non-imaging Measures of Health and Behavior

The HCP dataset provides comprehensive information about non-imaging measures (NIMs) regarding the participants’ physical and mental health, cognitive characteristics, and lifestyle. We excluded NIMs where >90% of the sample endorsed the same response, had >10% missing values, or were highly colinear (r >0.9) (Supplemental Table S1). Amongst the retained NIMs, we used age-adjusted measures when available and imputed missing values using the “mice” R package (Multivariate Imputation by Chained Equations; van Buuren & Groothuis-Oudshoorn, 2011). Based on prior literature (Chan et al., 2018; Elliott et al., 2019; Gurholt et al., 2021; Hatton et al., 2018; Karama et al., 2015; Steffener et al., 2016; Topiwala et al., 2017; Wertz et al., 2021), we distinguished between NIMs considered as factors that can potentially increase or decrease G-brainAGE (i.e., predictors) and those NIMs that can be considered functional correlates of G-brainAGE.

#### Predictors of G-brainAGE

NIMs considered as predictors of G-brainAGE pertained to (1) sociodemographic characteristics (e.g., sex, education); (2) quality of interpersonal relationships (e.g., loneliness, emotional support); (3) mental health (e.g. personal and parental psychopathology); (4) physical health (e.g. body mass index, blood pressure); (5) substance use (e.g., parental history and personal history of alcohol and substance use); and (6) female reproductive health (e.g. menstrual history and birth control use). In total, we considered 28 NIMs as predictors of interest for G-brainAGE for both sexes and an additional 4 NIMs pertaining to reproductive health for females only. Detailed definitions of these NIMs and the instruments used for their assessment are provided in supplemental material and Supplemental Table S2. Their distribution in the sample is shown in Supplemental Table S4.

#### Functional indicators of brain ageing

NIMs considered as functional indicators of brain ageing pertained to fluid and crystalized intelligence and physical fitness as captured by submaximal endurance, gait speed, and hand grip strength (Supplemental Table S3). These variables were chosen based on current consensus that they are reliable and sensitive measures of age-related frailty (Belsky et al., 2015; Kennedy et al., 2014). Their distribution in the sample is shown in Supplemental Table S4.

### Statistical Approach

#### Sex differences in G-brainAGE and L-brainAGE

Sex differences were computed on a subset of unrelated individuals (i.e., one randomly selected participant per family, n = 445). As G-brainAGE was already corrected for age, chronological age was entered as a covariate in the models for L-brainAGE. Statistical significance was set at P_FWE_ < .05 after family-wise error (FWE) correction.

#### Selection of predictors of brain ageing

To enhance interpretability and reduce the number of statistical tests we focused on potential predictors (Supplemental Table S2) that had at least a nominal association with G-brainAGE in this sample. These same predictors were then tested for their association with L-brainAGE. Although, some predictors may be associated with L-brainAGE alone, we selected to focus on those that seem to also have a global influence on brain-ageing as these are likely to be more meaningful for prevention and intervention strategies. Accordingly, and separately for each sex, we selected predictors that showed at least minimal univariate associations with G-brainAGE (rho > 0.1 or R^2^ > 0.01 for continuous and categorical variables, respectively).

#### Predictor importance for G-brainAGE

The selected predictors were entered into sex-specific random forest (RF) regression models to test their association with G-brainAGE (Breiman, 2001). RF regression is an ensemble machine learning method, which involves construction of multiple decision trees (i.e., forests) via bootstrap (bagging) and aggregates the predictions from these multiple trees to reduce the variance and improve the robustness and accuracy. For each bootstrapped sample, a portion of the observations (out-of-bag; OOB) are withheld and not used in the construction of the trees. RF allows for an importance measure to be determined for each predictor by measuring the effect of predictor permutation on the model’s mean decrease in accuracy. RF with 10-fold cross-validation was implemented using the “randomForest” package in R using 500 trees and a minimum terminal node sample size of 5. Importance values were scaled to range from 0-100 and then, for each predictor, averaged across folds. The directionality of relationships between the predictors and G-brainAGE were determined by examining beta values obtained by linear regression.

#### Brain-ageing predictors and L-brainAGE

The sex-specific predictors selected for the G-brainAGE analyses were entered into sex-specific, voxel-wise multiple regression models to examine their relationships with L-brainAGE, on the same subset used to test for sex differences in L-brainAGE. For both models, chronological age was entered as a covariate. Statistical significance was set at P_FWE_ < .05.

#### Functional indicators of brain-ageing

Associations between functional indicators of brain-ageing were determined for G-brainAGE only using Spearman’s correlations. To account for dependence between observations due to relatedness in the HCP data, stratified bootstrapping was carried out with 100 iterations such that each bootstrapped sample consisted of unrelated individuals (i.e., one randomly selected participant per family, n = 445).

#### Supplemental analyses for G-brainAGE

We used the procedures described above to identify predictors of G-brainAGE in the entire sample (i.e., including both sexes).

## Results

### Sex-differences in G-brainAGE and L-brainAGE

Model performance for global brain-age prediction was similar in females and males; the MAE was 2.72 years for both sexes and the respective R^2^ was 93% and 96%. There was no residual association between chronological age and G-brainAGE corrected for age-bias for either sex (Supplemental Figure S1). The mean and standard deviation (SD) of the uncorrected G-brainAGE was 0.03 (3.25) years for females and −0.07 (3.31) years for males; the difference amounts to approximately 1 calendar month and was not significant when age-corrected G-brainAGE values were used.

However, compared to males, females had significantly lower L-brainAGE in anterior brain regions, and specifically in the ventral and dorsal medial prefrontal cortex, the ventrolateral pre-frontal cortex, and the insula, and significantly higher L-brainAGE in posterior regions that included the cerebellum and brainstem (P_FWE_ < .05; Figure 1). Local MAE values showed a ventral to dorsal and a posterior to anterior gradient of decreasing sex differences (Supplemental Figure S2).

**Figure 1.**
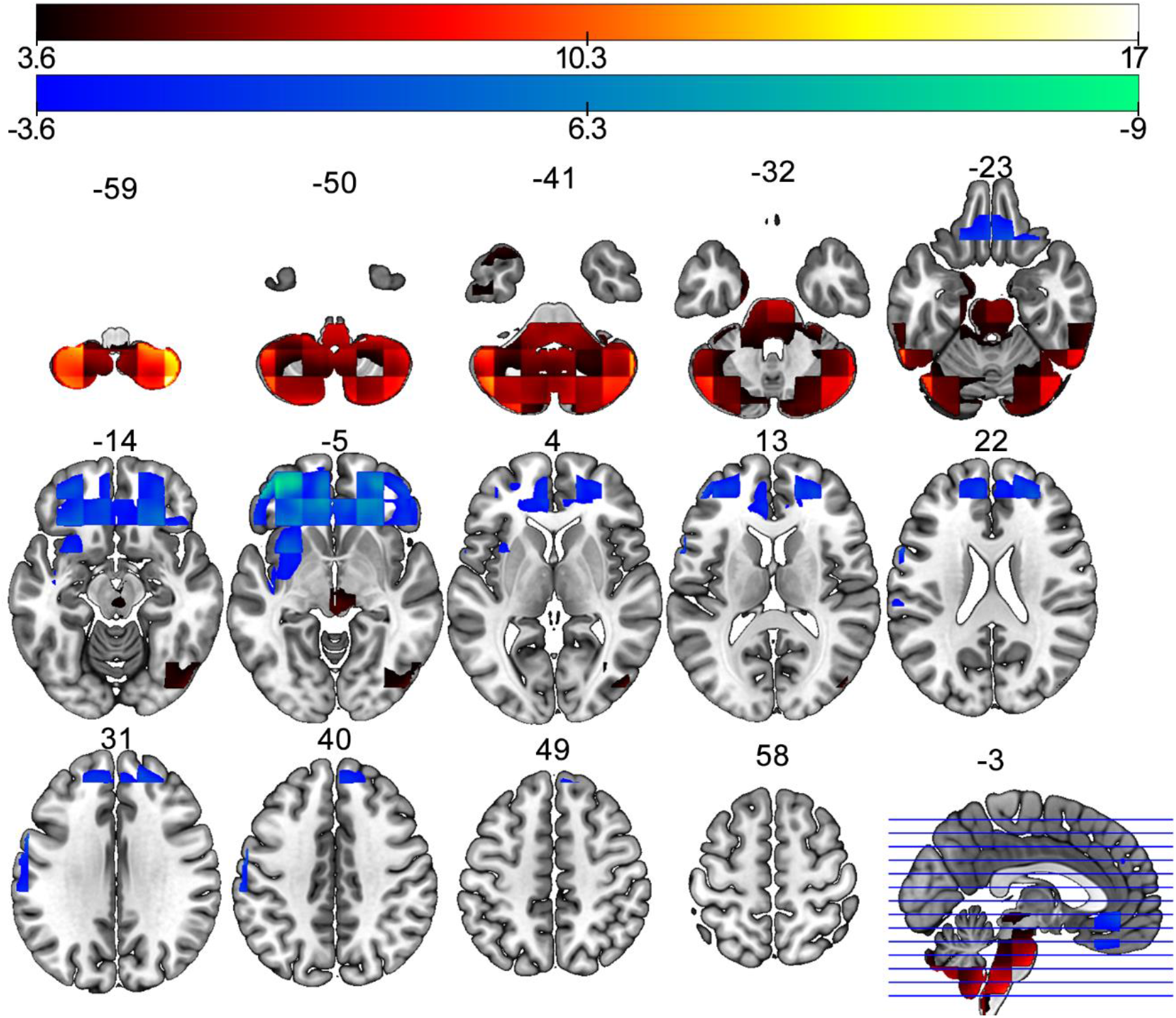
Sex differences in L-brainAGE. T-value overlay of statistically significant sex differences in L-brainAGE (P_FWE_ < .05 with familywise error-correction). Red/yellow: females > males; blue: males > females. Images are displayed in neurological orientation with MNI coordinates.

### Predictor Importance for G-brainAGE in Females and Males

Univariate associations with G-brainAGE identified largely different NIMs for females and males (Supplemental Table S5), which were then entered into sex-specific RF models. Accordingly, the RF model for females included 3 NIMs (i.e., systolic blood pressure, poor sleep quality and years of education; Figure 2A) and the corresponding model for males included 5 NIMs (i.e., race, poor sleep quality, childhood conduct problems, times used illicit drugs, and emotional support; Figure 2B). Figure 2 presents the importance of each predictor in the sex-specific models. Supplemental analyses including both sexes identified poor sleep quality, non-white race and time used illicit drugs as predictors of higher G-brainAGE (Supplemental Figure S3).

**Figure 2.**
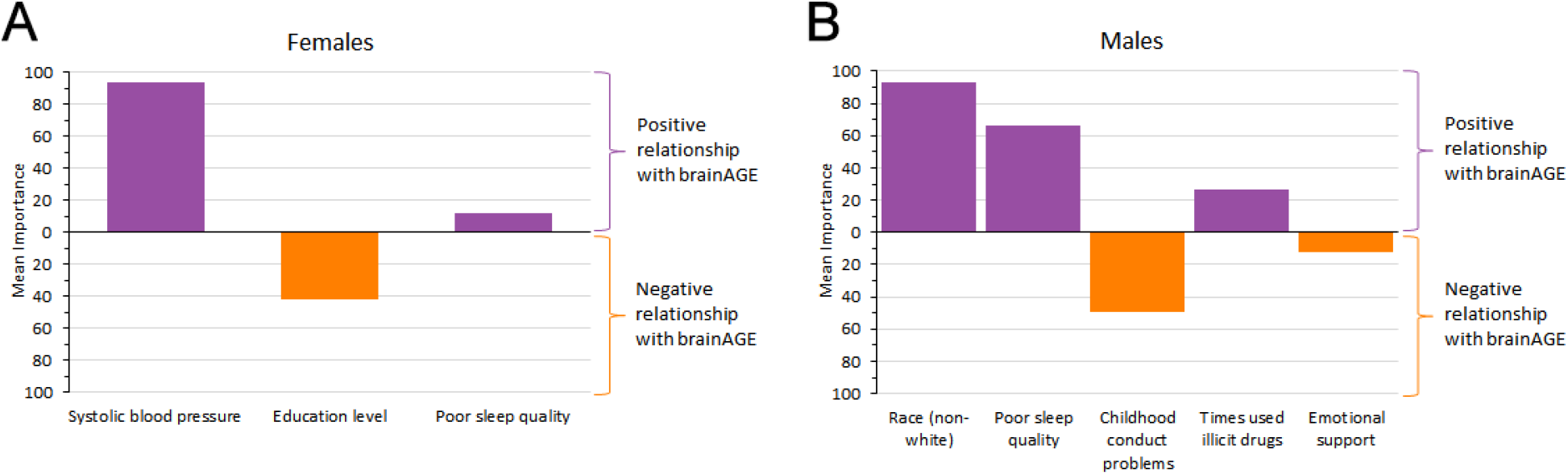
Predictors of G-brainAGE in females and males. Relative importance of predictors derived from random forest regression based on mean decrease in prediction accuracy when removed from model, scaled to range from 0-100; predictors are displayed in descending order of importance; (A) In females, there is a positive relationship with G-brainAGE for systolic blood pressure and poor sleep quality, and a negative relationship for education level; (B) In males, there was a positive relationship with G-brainAGE for non-white race, poor sleep quality, and times used illicit drugs, and a negative relationship for number of childhood conduct problems and emotional support.

### Voxel-level Associations Between L-brainAGE and Brain-ageing Predictors

Sex-specific, voxel-wise multiple regression models between L-brainAGE and the same predictors using in the preceding analyses for G-brainAGE did not identify significant associations in females at P_FWE_ < .05; at the uncorrected level, some positive associations were noted for systolic blood pressure (Supplemental Figure S4). In males, significant positive associations at P_FWE_ < .05 were found for black race and for poor sleep quality. The associations with Black race were widespread (Supplemental Figure S5A) while associations with poor sleep quality appeared mostly localized in the cerebellum (Supplemental Figure S5B).

### Functional Indicators of Brain-ageing

In both sexes, there were minimal and non-significant associations between G-brainAGE and endurance, gait speed, grip strength, composite score for fluid intelligence, and composite score for crystalized intelligence (Figure 3A-E and Supplemental Table S6).

**Figure 3.**
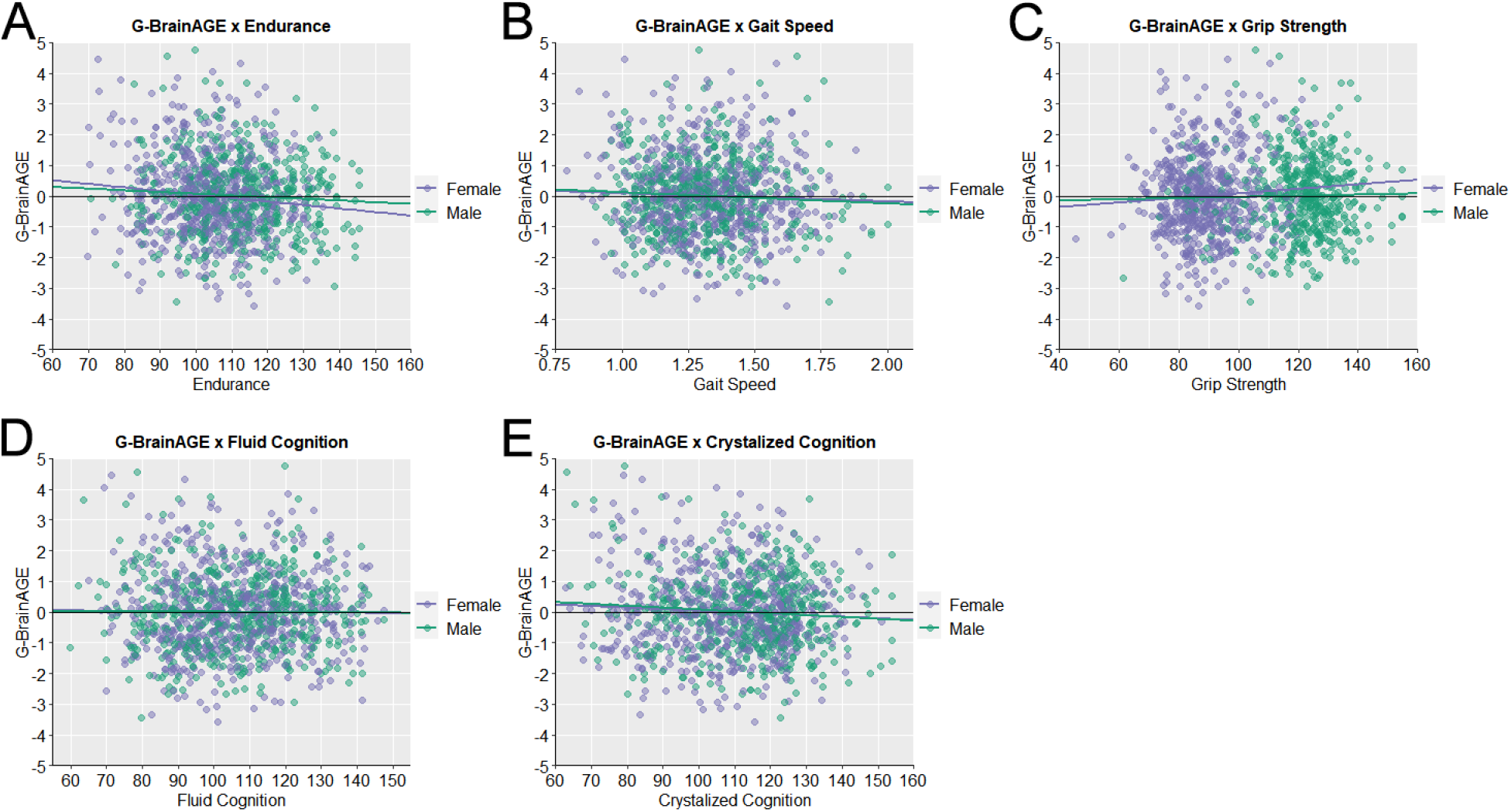
Scatter plots of the association of G-brainAGE and functional indicators. Results are shown for (A) endurance, (B) gait speed, (C) grip strength, (D) fluid cognition composite score, and (E) crystalized cognition composite score; green = males; purple = females. None of these associations were statistically significant.

## Discussion

The present study used machine learning methods to test for sex-differences in G-brainAGE and L-brainAGE and in the predictors of higher G-brainAGE in young adults. While G-brainAGE showed negligible sex-differences, L-brainAGE estimates indicated a more “youthful” appearing brain pattern in females compared to males in prefrontal cortical regions, while the opposite was the case for cerebellar regions. Poor sleep quality emerged as an important predictor of higher G-brainAGE in both men and women, while other predictors were sex-specific. Notably, non-white race was the most important contributor to higher G-brainAGE in males, while higher systolic blood pressure was the most important contributor to higher G-brainAGE in females.

### Sex-differences in Regional but not Global Ageing Patterns in Young Adults

G-brainAGE is a summary index of the global pattern of apparent brain structural ageing with positive and negative values being respectively indicative of accelerated or delayed ageing. In the current sample, the mean uncorrected G-brainAGE in both sexes was close to zero, suggesting that their global ageing pattern was congruent with their chronological age. These findings are aligned with prior studies demonstrating that sex-differences in G-brainAGE that emerge during adolescence appear to attenuate in early and middle adulthood (Bittner et al., 2021; Brouwer et al., 2021). Application of U-Net, a novel machine learning algorithm applied to brain structural data, enabled us to obtain regional estimates of brain-ageing, thus revealing fine-grained sex differences. The L-brainAGE findings suggest that female brains are more “youthful” than male brains in prefrontal cortical regions, while male brains are more youthful in cerebellar regions. Ventral cerebellar regions showed a higher degree of MAE in brain-age estimates than in prefrontal cortical regions, which may have a bearing on the results (Supplemental Figure S2). However, the pattern identified here closely mirrors the regional gradient of aerobic glycolysis in the human brain, whereby resting glucose consumption is about ten times higher in prefrontal regions compared to the cerebellum (Goyal et al., 2014; Vaishvani et al., 2010) This regional pattern parallels sex-differences in the expression of genes involved in neuronal integrity, protein synthesis and energy production (Berchtold et al., 2008; Goyal et al., 2019; Goyal et al., 2014; Skene et al., 2017). Expression of these genes is higher in females than males across adulthood (Berchtold et al., 2008; Skene et al., 2017) and has been associated with more youthful metabolic ageing patterns in females than in males (Goyal et al., 2019). Our results therefore suggest the possibility that the sex-differences in regional rate of age-related brain structural changes may reflect transcriptomic and metabolic mechanisms that could be examined further in future studies.

### Functional Association of G-brainAGE

We found no association between G-brainAGE with cognitive ability or measures of physical fitness, which is perhaps not unexpected since there was little evidence of accelerated global brain ageing in the current sample.

### Lower Sleep Quality as a Predictor of Higher G-brainAGE in Females and Males

Sleep is essential in maintaining homeostasis via multiple cellular, immune, and metabolic pathways (Zielinski, McKenna, & McCarley, 2016). Within the brain, sleep exerts powerful effects on molecular, cellular and network mechanisms of plasticity (Abel, Havekes, Saletin, & Walker, 2013). Even minor decrements in sleep quality disrupt circadian rhythms, impair the clearance of misfolded proteins, and induce molecular and cellular changes conducive to neuroinflammation and oxidative stress (reviewed by Bishir et al., 2020). The current findings reinforce this prior literature in demonstrating an association between poor sleep quality and higher G-brainAGE. Although a similar proportion of female (32.84%) and male (30.57%) participants reported sleep problems (Supplemental Table S4), poor sleep quality appeared to play a more important role for G-brainAGE and L-brainAGE in males. Compared to females, males have less slow-wave-sleep, which shows steeper age-related decline beginning in early adulthood (Yetton et al., 2018). Prior studies suggested an association between this age-related reduction in sleep quality with greater cortical thinning (Goldstone et al., 2018) which is captured by the G-brainAGE measure here. Additionally, the L-brainAGE analyses highlight the importance of cerebellar age-related reduction in males. These findings resonate with those of Zhou and colleagues (Zhou et al., 2017) who reported an association between increased brain-ageing in terms of cortico-cerebellar connectivity and poor sleep. However, we acknowledge that this explanation is only speculative since sleep architecture was not captured in the present study.

### Sex-specific Predictors of G-BrainAGE

The sex-specific analysis was more informative than when males and females were combined, underscoring the advantage of this approach. Numerically fewer predictors showed an association with G-brainAGE in females compared to males, with higher systolic blood pressure having the largest contribution to G-brainAGE and L-brainAGE in females but not in males. This observation accords with findings in a large longitudinal population-based sample (n = 7485; age range 20-76 years) where genetic risk factors and hypertension in early adulthood were the only predictors of late-life decline in females (Anstey et al., 2021). This relative paucity of potentially modifiable predictors of brain ageing in females in early adulthood could potentially contribute to their higher rates of dementia (Niu et al., 2017). We did not find significant associations between G-brainAGE and variables reflecting reproductive activity in the young females in this sample. Estrogens are considered neuroprotective (Gould et al., 1990; McCarthy, 2008) and may contribute to decelerate brain ageing in older females (Maki & Resnick, 2001). In the present study, sex hormone levels were not available, thus precluding direct assessment of their association with G-brainAGE. Nevertheless, it is possible that hormonal effects on brain morphology are less important in early adulthood but may be preconditions for maintaining a more youthful brain and/or might delay brain ageing processes much later in life. Longitudinal studies taking a lifespan perspective would be required to address these issues.

Educational level made a minor contribution to G-brainAGE in females but had no significant association with G-brainAGE in males. Although earlier research had suggested that educational attainment slows the rate of brain ageing (Steffener et al., 2016), this finding is consistent with the more recent evidence available that suggests that educational attainment has minimal, if any, influence on brain ageing (Nyberg et al., 2021).

In males, race emerged as the most important contributor to G-brainAGE and L-brainAGE; being non-white was associated with higher G-brainAGE. Prior studies have suggested that there is a greater burden of age-related disorders in middle-aged and older non-white individuals, particularly those of African ancestry, which has been considered indicative of greater age-related decline in brain organization (Amariglio et al., 2020; Gottesman et al., 2015). Biomarker-based indices of biological ageing also show a similar racial pattern (Levine & Crimmins, 2014). The advantage of the current study is that the sex-specific analyses undertaken highlight the importance of racial difference for males. The reasons for these racial differences in ageing remain poorly specified but are commonly attributed to the greater social and economic adversity experienced by non-white groups (Williams & Sternthal, 2010).

### Limitations

In addition to issues raised in previous sections, several strengths and limitations are worth further discussion. The study is cross-sectional and as such it does not address either causality or the longitudinal evolution of the reported findings. Participant sex was based on self-report and was not genetically determined; we consider the frequency of discrepancies between genetic versus reported sex to be too low to significantly influence the reported results. Sex, as defined here, incorporates societal and lifestyle differences that may differ between females and males. However, the sex-specific analyses performed in terms of predictors of G-brainAGE address this issue to some extent. The list of environmental exposures considered was substantial but not exhaustive. The HCP dataset does not include information on physical activity, which is considered protective based on its association with better cognitive performance in older adults (Colcombe & Kramer, 2003; Hughes et al., 2009), preserved gray matter (Colcombe et al., 2003), brain metabolic activity (Engeroff et al., 2019), and with lower brainAGE (Bittner et al., 2021). The analytical approach is a particular strength of this study as it combined two different machine learning methods to identify the regional age-related patterns of brain structure and the predictors of global age-related brain structure.

### Conclusions

The results presented here demonstrate the value of applying sex-specific analyses and machine learning methods to assess factors that influence the rate of age-related brain structural changes and their regional pattern. These findings might be useful in improving our understanding of sex-related differences in ageing and in identifying modifiable factors that influence the rate of age-related biological processes. Further investigations in longitudinal cohorts are needed to determine how sex and gender might affect the trajectory of human brain ageing.

