## Supplemental for "Sex differences in predictors and regional patterns of brain-age-gap estimates"

**Supplemental Information**

**A. Supplemental Methods ... 3**

**1. Neuroimaging data acquisition ... 3**

**2. Neuroimaging data processing for global and local brainAGE computation ... 3**

**3. Local brainAGE (L-brainAGE) computation ... 3**

**4. Global brainAGE (G-brainAGE) computation ... 5**

**5. Non-imaging variables ... 6**

Excluded variables. ... 6

Supplemental Table S1. Excluded non-imaging variables. ... 6

Predictors of interest for G-brainAGE. ... 7

Supplemental Table S2. Non-imaging HCP variables considered as predictors for G-brainAGE ... 8

Functional correlates of G-brainAGE. ... 10

Supplemental Table S3. Non-imaging HCP variables considered as functional correlates of G-brainAGE ... 10

**6. Distribution of non-imaging measures used as predictors and functional correlates of G-brainAGE in the study sample ... 10**

Supplemental Table S4. Descriptive statistics of the non-imaging measures used as predictors and functional correlates of G-brainAGE ... 10

**B. Supplemental Results ... 13**

**1. Correlations between G-brainAGE and chronological age ... 13**

Supplemental Figure S1. Correlation between chronological and G-brainAGE in females and males. ... 13

**2. Mean Absolute Error (MAE) in L-brainAGE prediction ... 13**

Supplemental Figure S2. Spatial distribution of MAE in L-brainAGE prediction. ... 14

**3. Univariate associations of predictors with G-brainAGE ... 14**

Supplemental Table S5. Univariate associations of predictors with G-brainAGE ... 14

### A. Supplemental Methods

#### 1. Neuroimaging data acquisition

The Human Connectome Project (HCP) data were acquired on a 3T Siemens Connectome-Skyra scanner using a 3D magnetization-prepared rapid gradient-echo (MPRAGE) sequence with the following parameters: repetition time (TR)/time to echo (TE)/inversion time (TI) = 2400/2.14/1000 ms, voxel size = 0.7 mm isotropic, flip angle = 8°, field of view (FOV) = 224 × 224 $mm^2$, duration of acquisition: 7 min 40 sec. Head movement artifacts were minimized by using strict criteria regarding head motion and overall quality control (Marcus et al., 2013); all data were de-identified prior to release (Van Essen & Barch, 2015).

#### 2. Neuroimaging data processing for global and local brainAGE computation

After downloading the data from the HCP repository, preprocessing was carried out using the standard pipelines implemented in the Statistical Parametric Mapping (SPM12) software package (https://www.fil.ion.ucl.ac.uk/spm/software/spm12/) to derive input features for global and local brain age prediction.

#### 3. Local brainAGE (L-brainAGE) computation

The local brain-age method applies deep learning to structural neuroimaging data to estimate “brain-age” at a local level. In contrast to global brain-age estimates, local estimates provide information about the spatial pattern of age-related brain changes. In this manuscript, we calculated local brain-age using the methods developed by Popescu, Glocker, Sharp, and Cole (2021); the full details of the framework are available at https://github.com/SebastianPopescu/U-NET-for-LocalBrainAge-prediction and are presented briefly below.

Popescu and colleagues collated data from multiple cohorts to create a discovery sample comprising a total of 3463 $T_1$-weighted MRI brain scans from healthy people (aged 18-90 years). An independent sample of 692 healthy people was used as the held-out test sample. All $T_1$-weighted brain MRI scans were pre-processed using the Statistical Parametric Mapping (SPM12) software package (https://www.fil.ion.ucl.ac.uk/spm/software/spm12/). Brain tissue was segmented into grey matter (GM) and white matter (WM) maps which were registered to the Montreal Neurological Institute 152 (MNI152) space using a nonlinear registration procedure implemented with the DARTEL algorithm (Ashburner, 2007), and were then resampled to 1.5$mm^3$ with a 4mm smoothing kernel. The local brain-age estimates were derived using a convolutional neural network adopted by the U-Net architecture introduced by Ronneberger, Fischer, and Brox (2015).

Data from the discovery sample (n = 3463) were randomly subsampled and split into a training (80%) and validation set (20%). The model parameters were determined in the training set and applied to the validation set. The training process involved entering the modulated GM and WM maps derived from SPM12 as input features to the network algorithm. The GM and WM maps images were split into overlapping 3-dimensional blocks of 523 voxels. The convolutional layers in the network used an isotropic 3×3×3 filter, convolved over the input image after which element-wise multiplication with the filter weights and subsequent summation was performed at each location. Subsequently, to allow for non-linear modelling, the obtained values were processed using an “activation function”; specifically, the activation function LeakyReLu with α=0.2 was used. LeakyReLu(α) was defined as: LeakyReLu(x) = max(x, 0) + min(x * α, 0), which allows a small, non-zero gradient when the unit is not “active”. The convolution operation was also controlled by its stride, which is how many voxels are skipped after every element-wise weight multiplication and summation. The value of stride was set to 1. The algorithm used down-sampling, which increases the effective field of view or “receptive field” of layers higher in the hierarchy. Down-sampling at each scale was implemented using two consecutive 3D 3×3×3 filter kernels with an initial number of channels set to 64, which was multiplied by 2 further down the down-sampling path. Down-sampling involved 2×2×2 average pooling. For the up-sampling part of the network, the down-sampling architecture was inverted by replacing the down-sampling layers with 2×2×2 up-sampling layers. Each convolution used a squeeze-and-excite unit based on the Squeeze & Excite networks (Hu, Shen, & Sun, 2018) to obtain age predictions over 123 voxels blocks. Voxel-level mean absolute error (MAE) cost function on the output layer and two additional cost functions at the two other scales of the architecture were used: global average pooling followed by a dense layer to predict brain-age at block-level. The model was implemented in TensorFlow (Abadi et al., 2016).

Neuroimaging-based age-prediction is subject to regression dilution leading to a greater under- or over-estimate of age, the further away a sample is from the training set mean age. To account for this effect, Popescu et al. used a separate small batch (n = 200) of participants randomly selected from the held-out dataset and obtained their voxel-level brain-age delta $\Delta_{i,v}$, (i.e., predicted minus actual age), where *i* indicates the *i*-th participant and *v* the *v*-th voxel. Then participants were grouped according to their chronological aged into 5-year bins, with the first bin covering participants with a chronological age between 18-25 years. For each bin *b*, the corresponding average voxel-level brain-age delta $\Delta_{b,v}$ was calculated, which represents the average brain-age delta for that voxel given the chronological age interval. Subsequently, to debias the voxel-level brain-age delta for a new participant (e.g., from testing set), $\Delta_{j,v}$, the following formula was used:

$$\Delta_{\tilde{j},v} = \Delta_{j,v} - \Delta_{b,v}$$

The accuracy of the model parameters from the discovery dataset was then tested in the held-out sample (n = 692). The voxel-level MAE (unadjusted) values of the model varied in different brain regions; lower values were in the prefrontal cortex and subcortical regions and higher in the occipital lobe, cerebellum, and brainstem.

**4. Global brainAGE (G-brainAGE) computation**

Downloaded $T_1$-weighted images of HCP participants were processed using standard pipelines implemented in SPM12 and the computational anatomy toolbox (CAT12) (Gaser & Dahnke, 2016; http://www.neuro.uni-jena.de/cat12/CAT12-Manual.pdf). Data from females and males were analysed separately but using identical procedures. Structural MRI data were preprocessed using the CAT12 toolbox (http://www.neuro.uni-jena.de/cat/), an extension of the SPM12 software (Wellcome Department of Cognitive Neurology, London, UK; http://www.fil.ion.ucl.ac.uk/spm/software/spm12/). The CAT12 toolbox extends the unified segmentation model into GM, WM and cerebrospinal fluid (CSF), executed through SPM12 (Ashburner and Friston, 2005) from Welcome Department of Clinical Neurology running on MATLAB2015a (The MathWorks, Natick, MA, USA) by (1) applying an Adaptive Maximum a Posterior (AMAP) technique, based on modeling local parameter variances as spatial functions (Rajapakse et al., 1997), (2) performing a Partial Volume Estimation (PVE), which estimates the ratio of pure tissue types in each voxel (Tohka et al., 2004) and (3) denoising using a classical Markov Random Field model (Rajapakse et al., 1997) for post-processing. A second denoising method employed after intensity normalization is a Spatial-Adaptive Non-Local Means (SANLM) filter (Manjón et al., 2010). The DARTEL algorithm (Ashburner, 2007) was used to normalize the GM and WM maps to the MNI (Montreal Neurological Institute) structural template. Final images were modulated with the Jacobian determinant generated through non-linear spatial normalization and lastly smoothed with a 10-mm Full-Width-at-Half-Maximum Gaussian kernel.

Support vector regression (SVR) implemented in NeuroMiner (https://github.com/neurominer-git/NeuroMiner-1) was used to predict age from each participant's GM volume map derived from the CAT12 toolbox.

In each model, the neuroimaging features were randomly split into 10 non-overlapping samples (outer loop). A 10-fold cross-validation was then performed on the nine folds constituting the training set whilst the 10th fold was held out as the test set. Within this inner loop, one of the 10 folds was again iteratively held out as a test set. This approach maximizes the available data by repeatedly splitting the dataset into training and test sets and uses the held-out test set to assess the model's performance whilst generating robust parameters that are resistant to overfitting. Feature-wise standardization (scaling between 0 and 1 and regressing out ICV) was applied during the inner cycle. Model accuracy was measured via the mean absolute error (MAE). Each test person's age was estimated by applying the model to the test set; age estimates were averaged across the models for which the subject in question was not part of the training set. G-BrainAGE was calculated in each participant as the difference between the predicted age and chronological age. G-BrainAGE estimates were corrected for any residual effects of age as per Le et al. (2018), and the corrected estimates were used in all subsequent analyses unless otherwise specified.

#### 5. Non-imaging variables

The non-imaging measures (NIMs) were ascertained using several study-specific questionnaires as well as validated instruments. Full details can also be accessed at: https://www.humanconnectome.org/documentation/Q3/HCP_Q3_Release_Appendix_VII.pdf.

***Excluded variables*.** In the first instance, we excluded NIMs of potential interest where >90% of the sample endorsed the same response, had >10% of missing values, or were highly colinear (r >0.9). The excluded measures are shown in Supplemental Table S1.

**Supplemental Table S1. Excluded non-imaging variables.**

| Variable | Name of variable in the HCP database | Reason for exclusion |
|---|---|---|
| Thyroid stimulating hormone level | ThyroidHormone | 30% missing |
| Percentage of hemoglobin that is glycated | HbA1C | 31% missing |
| Parental history of schizophrenia/psychosis | FamHist_Moth_Scz and FamHist_Fath_Scz | 99.5% endorsed "no" |
| Parental history of bipolar disorder | FamHist_Moth_BP and FamHist_Fath_BP | 96.8% endorsed "no" |
| Parental history of anxiety | FamHist_Moth_Anx and FamHist_Fath_Anx | 93.4% endorsed "no" |
| Parental history of Alzheimer's Disease | FamHist_Moth_Alz and FamHist_Fath_Alz | 98.6% endorsed "no" |
| Parental history of Parkinson's Disease | FamHist_Moth_PD and FamHist_Fath_PD | 99.1% endorsed "no" |
| Parental history of Tourette's Syndrome | FamHist_Moth_TS and FamHist_Fath_TS | 100% endorsed "no" |

| Supplemental Table S1. Excluded non-imaging variables. | | |
|---|---|---|
| **Variable** | **Name of variable in the HCP database** | **Reason for exclusion** |
| No parental history of any neuropsychiatric disorder | FamHist_Moth_None and FamHist_Fath_None | 94.3% endorsed “yes” |
| Lifetime history of panic disorder | SSAGA_PanicDisorder | 92.7% endorsed “no” |
| Lifetime history of agoraphobia | SSAGA_Agoraphobia | 93.2% endorsed “no” |
| Lifetime history of diagnosed DSM IV major depressive episode | SSAGA_Depressive_Ep | 90.5% endorsed “no” |
| Lifetime number of DSM IV depressive symptoms | SSAGA_Depressive_Sx | Captured by prior variables |
| Met DSM IV criteria for Alcohol Dependence over lifetime | SSAGA_Alc_D4_Dp_Dx | 94.3% endorsed “no” |
| Number of DSM IV Alcohol Dependence criteria met | SSAGA_Alc_D4_Dp_Sx | As above |
| Number of symptoms of DSM IV Alcohol Abuse over lifetime | SSAGA_Alc_D4_Ab_Sx | As above |
| Times used cocaine | SSAGA_Times_Used_Cocaine | 92.9% “endorsed never” |
| Times used hallucinogens | SSAGA_Times_Used_Hallucinogens | Captured by “times used all classes of non-marijuana illicit drugs” |
| Times used opiates | SSAGA_Times_Used_Opiates | 91.5% endorsed “never” |
| Times used sedatives | SSAGA_Times_Used_Sedatives | 93.5% endorsed “never” |
| Times used stimulants | SSAGA_Times_Used_Stimulants | 90.5% endorsed “never” |
| Met DSM IV criteria for Marijuana Dependence over lifetime | SSAGA_Mj_Ab_Dep | 90.8% endorsed “no” |
| Colour vision | Color_Vision | 97.7% endorsed “normal” |
| History of hypothyroidism | Hypothyroidism | 99.7% endorsed “no” |
| History of hyperthyroidism | Hyperthyroidism | 99.7% endorsed “no” |
| History of other endocrine disorder | OtherEndocrn_Prob | 97.2% endorsed “no” |
| Current smoking status | SSAGA_TB_Still_Smoking | Colinear with # of days used tobacco in past 7 days |
| Ever used marijuana | SSAGA_Mj_Use | Colinear with “times used marijuana” |
| Smoking history | SSAGA_TB_Smoking_History | Captured by prior variables |
| Early Childhood Cognition Composite Score | CogEarlyComp_AgeAdj | Colinear with Fluid Cognition Composite Score. |

***Predictors of interest for G-brainAGE.*** NIMs considered predictors of G-brainAGE are shown in Supplemental Table S2. Most socioeconomic variables were acquired using study specific questionnaires. The Pittsburgh Sleep Quality Index (PSQI; Buysse, Reynolds, Monk, Berman, & Kupfer, 1989) and the Adult Self Report instrument (ASR 18/59; Achenbach, Ivanova, & Rescorla, 2017) were respectively used to assess sleep and internalizing and externalizing psychopathology. Interpersonal relationships were evaluated using instruments from the NIH toolbox (https://www.healthmeasures.net/explore-measurement-systems/nih-toolbox). Participants' height and weight measurements were used to calculate their individual Body Mass Index (BMI) (https://www.cdc.gov/healthyweight/assessing/bmi/adult_bmi/english_bmi_calculator/bmi_calculator.html) according to the United States Centers for Disease Control and Prevention; individuals with a BMI lower than 18.5 are considered underweight, those with a BMI between 18.5 and 24.9 are considered a healthy weight, those with a BMI between 25 and 29.9 are considered overweight, and a BMI of 30 and above indicates obesity. Participants' blood pressure (BP) was categorized based on their hypertension status according to the International Society of Hypertension Global Hypertension Practice Guidelines (Unger et al., 2020) as normal BP: <130 mmHg systolic BP (SBP); high-normal SBP: 130-139 mmHg SBP; Grade 1 hypertension: 140-159 mmHg SBP; and Grade 2 hypertension: ≥160 mmHg SBP.

**Supplemental Table S2. Non-imaging HCP variables considered as predictors for G-brainAGE**

| Domain | Name of variable in the HCP database | Definition and assessment |
|---|---|---|
| **Sociodemographic Characteristics** | | |
| Sex | Gender | Male; female |
| Race | Race | Self-reported racial identity (White; Black/African American; Asian/Native Hawaiian/Other Pacific Islander; mixed/other) |
| Employment | SSAGA_Employ | Employment Status (not working; working part-time; working full-time) |
| Income | SSAGA_Income | Annual income (low: < $40k; middle: $40k-$99,999k; high, >$100k) |
| Education | SSAGA_Educ | Years of education (12 years or less; 13-16 years; 17 years or more) |
| **Interpersonal Relationships** | | |
| Relationship Status | SSAGA_Rlshp | In a married/live-in relationship (no; yes) |
| Life Satisfaction | LifeSatisf_Unadj | NIH Toolbox: General Life Satisfaction Survey |
| Meaning and Purpose | MeanPurp_Unadj | NIH Toolbox: Meaning and Purpose Survey |
| Perceived Stress | PercStress_Unadj | NIH Toolbox: Perceived Stress Survey |
| Self Efficacy | SelfEff_Unadj | NIH Toolbox: Self-Efficacy Survey |
| Emotional Support | EmotSupp_Unadj | NIH Toolbox: Emotional Support Survey |
| Instrumental Support | InstruSupp_Unadj | NIH Toolbox: Instrumental Support Survey |
| Loneliness | Loneliness_Unadj | NIH Toolbox: Loneliness Survey |
| Friendships | Friendship_Unadj | NIH Toolbox: Friendship Survey |

| **Supplemental Table S2. Non-imaging HCP variables considered as predictors for G-brainAGE** | | |
|---|---|---|
| **Domain** | **Name of variable in the HCP database** | **Definition and assessment** |
| Perceived Hostility | PercHostil_Unadj | NIH Toolbox: Perceived Hostility Survey |
| Perceived Rejection | PercReject_Unadj | NIH Toolbox: Perceived Rejection Survey |
| **Mental Health** | | |
| ASR Internalizing Sum | ASR_Intn_T | Adult Self-Report (ASR): Internalizing T-score |
| ASR Externalizing Sum | ASR_Extn_T | Adult Self-Report (ASR): Externalizing T-score |
| Sleep Quality | PSQI_Score | Pittsburgh Sleep Quality Index (PSQI) Total Score |
| Childhood Conduct Problems | SSAGA_ChildhoodConduct | History of Childhood Conduct Problems (no history of problems; history of 1 problem; history of 2+ problems) |
| Parental History of Depression | FamHist_Moth_Dep and FamHist_Fath_Dep (variables combined to create single index of parental history) | Parental history of depression (no; yes) |
| **Physical Health** | | |
| Body Mass Index | BMI | Body mass index |
| Hematocrit | Hematocrit_1 | Hematocrit sample 1 |
| Systolic Blood Pressure | BPSystolic | Systolic blood pressure |
| **Substance Use** | | |
| Parental History of Drug/Alcohol Problems | FamHist_Moth_DrgAlc and FamHist_Fath_DrgAlc | Variables on maternal and paternal drug/alcohol problems were combined to create single binarized variable of parental history: Parental history of drug/alcohol problems (no; yes) |
| Alcohol use in past 7 days | Num_Days_Drank_7days | Number of days drank alcohol in past 7 days |
| DSM-IV Alcohol Abuse | SSAGA_Alc_D4_Ab_Dx | Meets/met DSM-IV criteria for Alcohol Abuse over lifetime (no; yes) |
| Tobacco use in past 7 days | Num_Days_Used_Any_Tobacco_7days | Number of days smoked/used any tobacco in past 7 days |
| Illicit drugs use ever | SSAGA_Times_Used_Illicits | Number of times used non-marijuana illicit drugs (never; 1-10 times; 11+ times) |
| Marijuana use ever | SSAGA_Mj_Times_Used | Number of times used marijuana (never; 1-10 times; 11-100 times; 100+ times |
| **Female Reproductive Health** | | |
| Age at first menstrual cycle | Menstrual_AgeBegan | Females only: Age at first menstrual cycle |
| Regularity of menstrual cycles | Menstrual_RegCycles | Females only: Regular menstrual cycles (no; yes) |
| Days since last menstrual period | Menstrual_DaysSinceLast | Females only: Number of days since last menstrual period occurred |
| Use of birth control drugs | Menstrual_UsingBirthControl | Females only: Use of birth control drugs (no; yes); No participant reported using fertility drugs |

***Functional correlates of G-brainAGE.*** All HCP participants underwent detailed cognitive assessment using the NIH Toolbox Cognition Battery that comprises tests of executive function, episodic memory, language, processing speed, working memory, and attention (Weintraub et al., 2013). In this paper, we use the composite scores for fluid and crystalized intelligence. Endurance was assessed by asking participants to walk as far as possible over 2 minutes. Participants undertook two trials and the longest distance walked in either trial was used as the outcome. Gait speed was assessed by asking participants to walk at a self-selected and comfortable pace over a straight distance of 6 meters, with the central 4 meters representing the testing zone. The outcome is the time taken to walk the 4 meters of the testing zone. Full force grip strength was measured with both hands using a Jamar Plus Digital dynamometer with the elbow bent to 90 degrees and arm against the trunk. The outcome measure was pounds of force for the dominant hand. Grip strength is strongly correlated with knee extension performance and is thus considered a measure of body strength (Bohannon, Magasi, Bubela, Wang, & Gershon, 2012). Details of the variables used as functional correlates of G-brainAGE are shown in Supplemental Table S3.

**Supplemental Table S3. Non-imaging HCP variables considered as functional correlates of G-brainAGE**

| **Domain** | **Name of variable in the HCP database** | **Definition and assessment** |
|---|---|---|
| Endurance | Endurance_AgeAdj | Physical Endurance (2-minute walk endurance test, age-adjusted score) |
| Gait Speed | GaitSpeed_Comp | 4-Meter Walk Gait Speed Rest |
| Grip Strength | Strength_AgeAdj | Grip strength test, age-adjusted score |
| Fluid Cognition Composite Score | CogFluidComp_AgeAdj | Fluid Cognition Composite, age-adjusted score |
| Crystalized Cognition Composite Score | CogCrystalComp_AgeAdj | Crystalized Cognition Composite, age-adjusted score |

#### 6. Distribution of non-imaging measures used as predictors and functional correlates of G-brainAGE in the study sample

Supplemental Table S4 shows the distribution of the non-imaging measures used as predictors or functional correlates of G-brainAGE in the study sample.

**Supplemental Table S4. Descriptive statistics of the non-imaging measures used as predictors and functional correlates of G-brainAGE**

| **Measure** | **Females** | **Males** |
|---|---|---|
| **Sociodemographic** | | |
| Race: White, N (%) | 451 (74.42%) | 398 (78.50%) |
| Race: Black or African American, N (%) | 103 (17.00%) | 65 (12.82%) |

| Supplemental Table S4. Descriptive statistics of the non-imaging measures used as predictors and functional correlates of G-brainAGE | | |
|---|---|---|
| **Measure** | **Females** | **Males** |
| Race: Asian/Native Hawaiian/Other Pacific Islander, N (%) | 35 (5.78%) | 30 (5.92%) |
| Race: Mixed/Other, N (%) | 17 (2.81%) | 14 (2.76%) |
| Employment: Unemployed, N (%) | 107 (17.66%) | 56 (11.05%) |
| Employment: Part-time, N (%) | 121 (19.97%) | 72 (14.20%) |
| Employment: Full-time, N (%) | 378 (62.38%) | 379 (74.75%) |
| Household Income: Low (< $40k), N (%) | 233 (38.45%) | 211 (41.62%) |
| Household Income: Mid ($40k - $99,999k), N (%) | 281 (46.37%) | 216 (42.60%) |
| Household Income: High ($100k+), N (%) | 92 (15.18%) | 80 (15.78%) |
| Education: < 13 years, N (%) | 104 (17.16%) | 91 (17.95%) |
| Education: 13-16 years, N (%) | 390 (64.36%) | 354 (69.82%) |
| Education: 17+ years, N (%) | 112 (18.48%) | 62 (12.23%) |
| **Interpersonal Relationships** | | |
| Relationship Status: Married/Live-in, N (%) | 305 (50.33%) | 187 (36.88%) |
| Life Satisfaction, Mean (SD) | 54.80 (9.47) | 54.22 (8.81) |
| Meaning and Purpose, Mean (SD) | 52.47 (8.64) | 50.94 (8.70) |
| Perceived Stress, Mean (SD) | 48.74 (8.98) | 47.86 (9.35) |
| Self Efficacy, Mean (SD) | 50.12 (7.99) | 51.98 (8.51) |
| Emotional Support, Mean (SD) | 52.48 (9.11) | 50.12 (9.91) |
| Instrumental Support, Mean (SD) | 48.18 (8.69) | 47.85 (9.37) |
| Loneliness, Mean (SD) | 51.08 (8.27) | 51.00 (9.01) |
| Friendships, Mean (SD) | 50.37 (9.17) | 50.42 (8.83) |
| Perceived Hostility, Mean (SD) | 47.65 (8.71) | 50.01 (8.33) |
| Perceived Rejection, Mean (SD) | 48.08 (8.71) | 48.99 (8.82) |
| **Mental Health** | | |
| Adult Self-Report Internalizing, Mean (SD) | 48.04 (10.17) | 49.02 (11.15) |
| Adult Self-Report Externalizing, Mean (SD) | 47.20 (8.91) | 50.55 (8.57) |
| Pittsburgh Sleep Questionnaire (PSQI), Mean (SD) | 4.94 (2.93) | 4.64 (2.53) |
| No sleep problems (PSQI<6), N (%) | 407 (67.16%) | 352 (69.43%) |
| Childhood Conduct Problems: None, N (%) | 433 (71.45%) | 241 (47.53%) |
| Childhood Conduct Problems: 1, N (%) | 130 (21.45%) | 179 (35.31%) |
| Childhood Conduct Problems: 2+, N (%) | 43 (7.10%) | 87 (17.16%) |
| Parental History of Depression, N (%) | 106 (17.49%) | 61 (12.03%) |
| **Physical Health** | | |
| Body Mass Index, Mean (SD) | 26.27 (5.78) | 26.84 (4.39) |
| Underweight, N (%)[1] | 21 (3.47%) | 10 (1.97%) |
| Healthy weight, N (%)[1] | 286 (47.19%) | 184 (36.29%) |
| Overweight, N (%)[1] | 159 (26.24%) | 210 (41.42%) |
| Obese, N (%)[1] | 140 (23.10%) | 103 (20.32%) |
| Hematocrit, Mean (SD) | 41.16 (4.11) | 45.94 (3.77) |
| Systolic Blood Pressure (SBP), Mean (SD)[2] | 119.71 (13.61) | 128.32 (13.30) |

**Supplemental Table S4. Descriptive statistics of the non-imaging measures used as predictors and functional correlates of G-brainAGE**

| Measure | Females | Males |
|---|---|---|
| Normal SBP, N (%)[2] | 326 (53.80%) | 129 (25.44%) |
| High-normal SBP, N (%)[2] | 157 (25.91%) | 155 (30.57%) |
| Grade 1 Hypertension, N (%)[2] | 77 (12.71%) | 123 (24.26%) |
| Grade 2 Hypertension, N (%)[2] | 46 (7.59%) | 100 (19.72%) |
| **Substance Use** | | |
| Parental History of Drug/Alcohol Problems, N (%) | 100 (16.50%) | 66 (13.02%) |
| Alcohol use in past 7 days, Mean (SD) | 1.41 (1.59) | 1.97 (1.96) |
| History of DSM-IV Alcohol Abuse, N (%) | 66 (10.89%) | 102 (20.12%) |
| Tobacco use in past 7 days, Mean (SD) | 0.69 (1.99) | 1.38 (2.59) |
| Non-marijuana illicit drug use: Never, N (%) | 506 (83.50%) | 367 (72.39%) |
| Non-marijuana illicit drug use: 1-10 times, N (%) | 70 (11.55%) | 74 (14.60%) |
| Non-marijuana illicit drug use: 11+ times, N (%) | 30 (4.95%) | 66 (13.02%) |
| Marijuana use: Never, N (%) | 303 (50.00%) | 202 (39.84%) |
| Marijuana use: 1-10 times, N (%) | 186 (30.69%) | 122 (24.06%) |
| Marijuana use: 11-100 times, N (%) | 61 (10.07%) | 67 (13.21%) |
| Marijuana use: 101+ times, N (%) | 56 (9.24%) | 116 (22.88%) |
| **Female Reproductive Health** | | |
| Age at first menstrual cycle, Mean (SD) | 12.70 (1.64) | - |
| Has regular menstrual cycles, N (%) | 477 (78.71%) | - |
| Days since last menstrual period, Mean (SD) | 136.16 (585.14) | - |
| Uses birth control drugs, N (%) | 172 (28.38%) | - |
| **Functional Correlates** | | |
| Endurance, Mean (SD) | 104.00 (12.52) | 112.12 (14.37) |
| Gait Speed, Mean (SD) | 1.32 (0.19) | 1.31 (0.20) |
| Grip Strength, Mean (SD) | 89.69 (11.88) | 120.14 (14.49) |
| Fluid Cognition Composite Score, Mean (SD) | 105.30 (16.57) | 105.77 (17.69) |
| Crystalized Cognition Composite Score, Mean (SD) | 107.66 (16.90) | 112.41 (17.08) |

[1]Status assigned based on body mass index.
[2]Blood pressure status assigned according to the International Society of Hypertension Global Hypertension Practice Guidelines.

#### B. Supplemental Results

##### 1. Correlations between G-brainAGE and chronological age

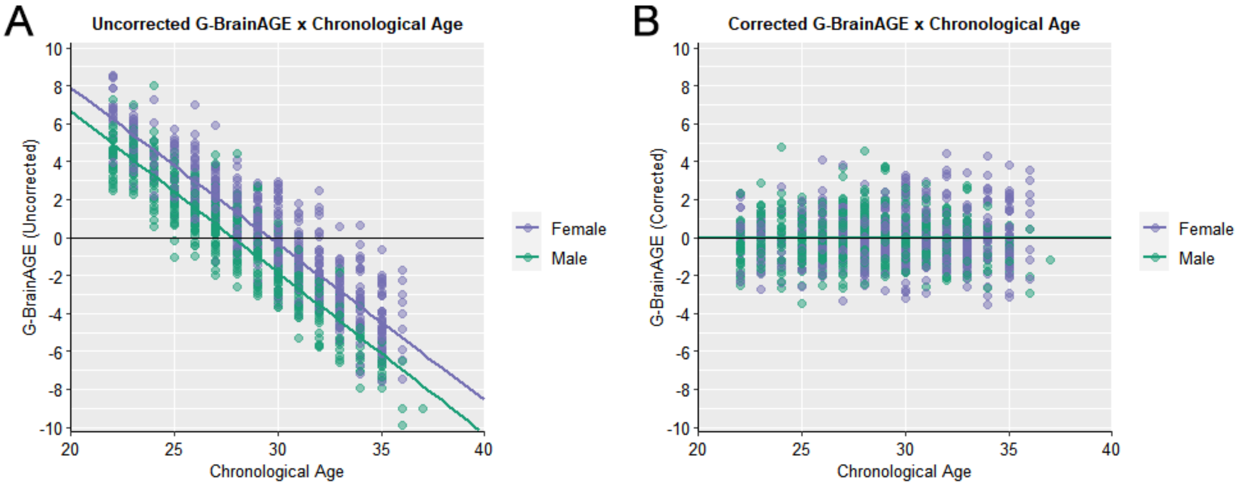

**Supplemental Figure S1. Correlation between chronological and G-brainAGE in females and males.** (A) Scatter plot illustrating negative correlations between uncorrected G-brainAGE and chronological age. (B) Scatter plot illustrating absence of residual associations between corrected G-brainAGE and chronological age.

##### 2. Mean Absolute Error (MAE) in L-brainAGE prediction

Voxel-wise local MAEs were computed in the same sample of participants as selected for examination of sex differences in L-brainAGE (i.e., one participant per family, n = 445). The spatial pattern showed a ventral to dorsal gradient of decreasing MAE (Supplemental Figure S2).

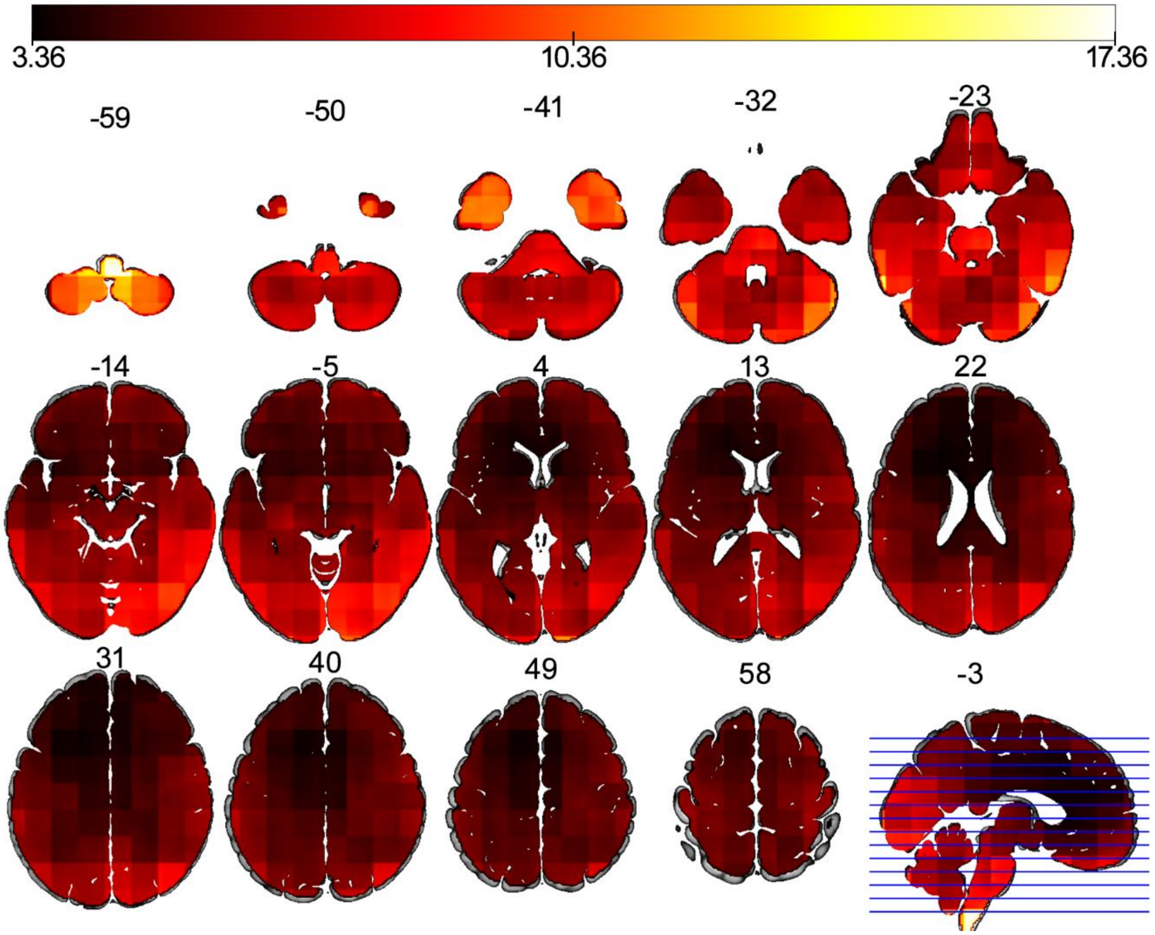

**Supplemental Figure S2. Spatial distribution of MAE in L-brainAGE prediction.** Spatial distribution of local mean absolute error (MAE) showing a ventral to dorsal and posterior to anterior gradient of decreasing MAE.

#### 3. Univariate associations of predictors with G-brainAGE

| **Supplemental Table S5. Univariate associations of predictors with G-brainAGE** | | | |
|---|---|---|---|
| | **Associations of G-brainAGE with predictor variables** | | |
| | **Females** | **Males** | **All** |
| **Continuous Variables** | | | |
| Life Satisfaction | -.03 (-.04, -.03) | -.07 (-.08, -.07) | -.06 (-.07, -.05) |
| Meaning and Purpose | -.01 (-.02, -.01) | -.08 (-.09, -.08) | -.05 (-.06, -.04) |
| Perceived Stress | -.04 (-.04, -.03) | .04 (.03, .04) | .01 (.00, .01) |
| Self-Efficacy | -.03 (-.03, -.02) | -.10 (-.10, -.09) | -.07 (-.07, -.06) |
| Emotional Support | .02 (.01, .02) | **-.12 (-.13, -.12)** | -.05 (-.05, -.04) |
| Instrumental Support | .02 (.02, .03) | -.06 (-.07, -.06) | -.03 (-.03, -.02) |
| Loneliness | -.01 (-.02, .00) | .06 (.06, .07) | .01 (.00, .01) |
| Friendships | .02 (.01, .03) | -.08 (-.08, -.07) | -.02 (-.03, -.02) |
| Perceived Hostility | .02 (.01, .02) | -.04 (-.05, -.03) | .00 (.00, .01) |

| **Supplemental Table S5. Univariate associations of predictors with G-brainAGE** | | | |
|---|---|---|---|
| | **Associations of G-brainAGE with predictor variables** | | |
| | **Females** | **Males** | **All** |
| Perceived Rejection | .01 (.01, .02) | .04 (.04, .05) | .02 (.02, .03) |
| ASR Internalizing Sum | -.06 (-.06, -.05) | .09 (.08, .09) | -.02 (-.03, -.01) |
| ASR Externalizing Sum | .03 (.02, .03) | .04 (.03, .04) | .02 (.02, .03) |
| Sleep Quality | **.13 (.13, .14)** | **.11 (.10, .11)** | **.12 (.11, .12)** |
| BMI | .02 (.02, .03) | .04 (.03, .04) | .04 (.03, .04) |
| Hematocrit | .05 (.04, .06) | -.03 (-.04, -.02) | .00 (-.01, .00) |
| BP Systolic | **.10 (.10, .11)** | .02 (.01, .02) | .09 (.09, .10) |
| No. of days drank alcohol (past 7 days) | .05 (.04, .06) | -.06 (-.07, -.05) | .00 (-.01, .00) |
| No. of days used tobacco (past 7 days) | .06 (.06, .07) | .06 (.05, .06) | .04 (.03, .05) |
| Age of first menstrual period | .01 (.00, .01) | - | - |
| No. of days since last menstrual period | .08 (.07, .08) | - | - |
| **Categorical Variables** | | | |
| Race | .006 (.005, .007) | **.032 (.030, .034)** | **.014 (.013, .016)** |
| Employment Status | .004 (.003, .004) | .002 (.002, .002) | .004 (.003, .005) |
| Household Income | .005 (.004, .006) | .005 (.004, .006) | .003 (.003, .004) |
| Education Level | **.024 (.023, .026)** | .002 (.002, .003) | .006 (.005, .007) |
| Relationship Status | .007 (.006, .008) | .007 (.006, .008) | .005 (.005, .006) |
| History of Childhood Conduct Problems | .005 (.004, .006) | **.024 (.022, .025)** | .010 (.008, .011) |
| Parental History of Depression | .003 (.003, .004) | .005 (.004, .005) | .001 (.001, .001) |
| Parental History of Drug/Alcohol Problems | .002 (.001, .002) | .007 (.006, .008) | .004 (.003, .004) |
| History of DSM-IV Alcohol Abuse | .001 (.001, .001) | .001 (.001, .001) | .001 (.001, .002) |
| Times used non-marijuana illicit drugs | .010 (.009, .012) | **.015 (.013, .016)** | **.012 (.011, .013)** |
| Times used marijuana | .010 (.009, .011) | .005 (.005, .006) | .006 (.005, .007) |
| Menstrual Regularity | .003 (.003, .004) | - | - |
| Birth Control Usage | .001 (.001, .001) | - | - |
| Associations were calculated using Spearman rank correlations for continuous variables and linear regression models for categorical variables. Stratified bootstrapping was performed to account for the inclusion of family members in the HCP data (i.e., only unrelated individuals were selected on each iteration). For continuous variables, the reported statistic is mean rho value and 95% confidence intervals. For categorical variables, the reported statistic is mean $R^2$ value and 95% confidence intervals. Values for predictors that were selected for random forest model inclusion are presented in bold font. | | | |

#### 4. Predictors of G-brainAGE in the entire sample

In addition to the sex-specific analyses in the main manuscript, we tested the significance of predictors of G-brainAGE in the entire sample (females and males together). NIMs with at least minimal association with G-brainAGE in the full sample (Supplemental Table S5) were entered as predictor variables in a single RF model that included both sexes together. Model implementation and predictor importance followed the same procedures described in the main text. The results are shown in Supplemental Figure S3; poor sleep quality, non-white race, and times used non-marijuana illicit drugs all had positive relationships with G-brainAGE.

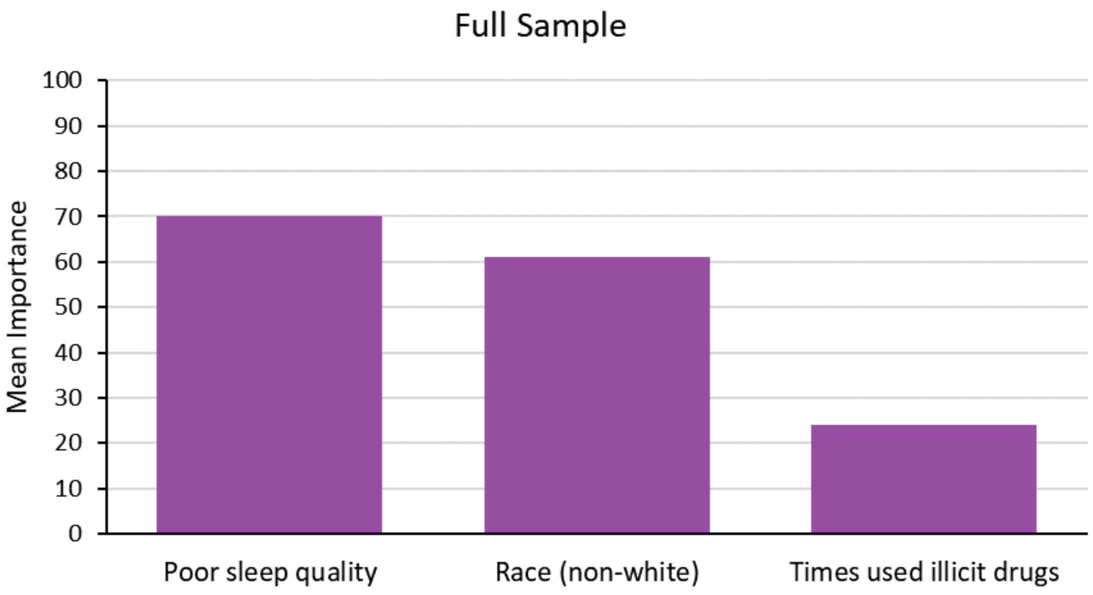

**Supplemental Figure S3. Predictor importance for G-brainAGE in the whole sample.**

**5. Voxel-wise association between key predictors of brain-ageing with L-brainAGE**

Predictors previously identified as important in the sex-specific random forest models were entered into sex-specific, voxel-wise multiple regression models to examine their relationships with L-brainAGE with chronological age as a covariate. We used the same subset of unrelated individuals that had been selected for examining sex differences in L-brainAGE. Each predictor was examined for significant main effects.

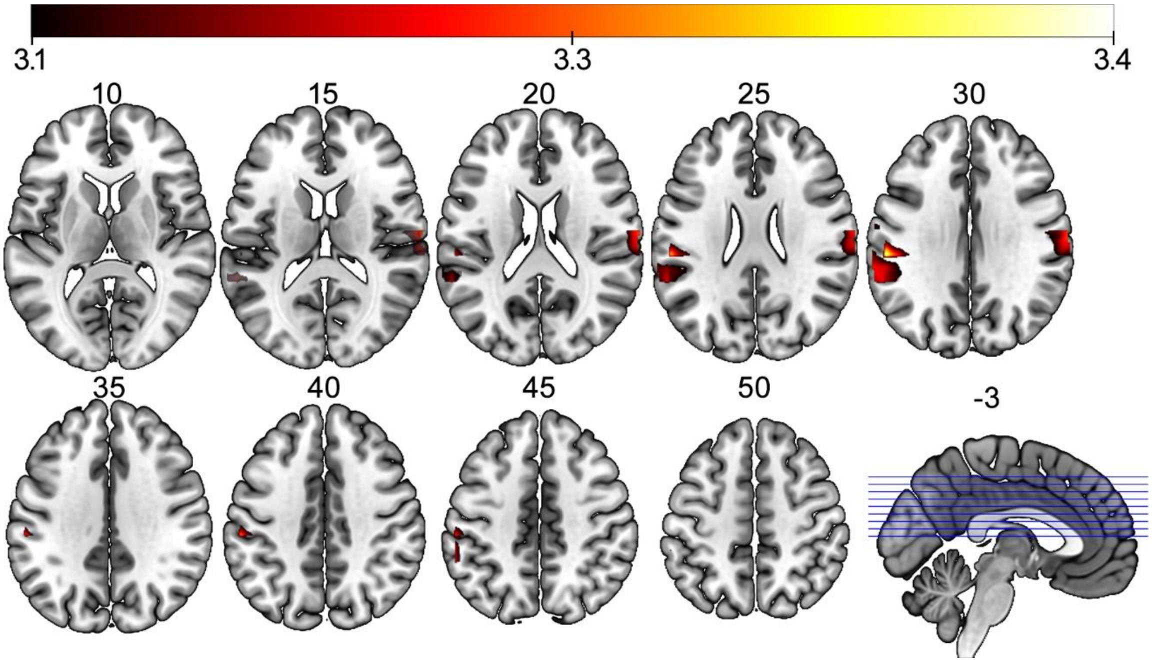

**Supplemental Figure S4. Voxel-wise association between systolic blood pressure and L-brainAGE in females.** Results are significant at uncorrected $P < .001$.

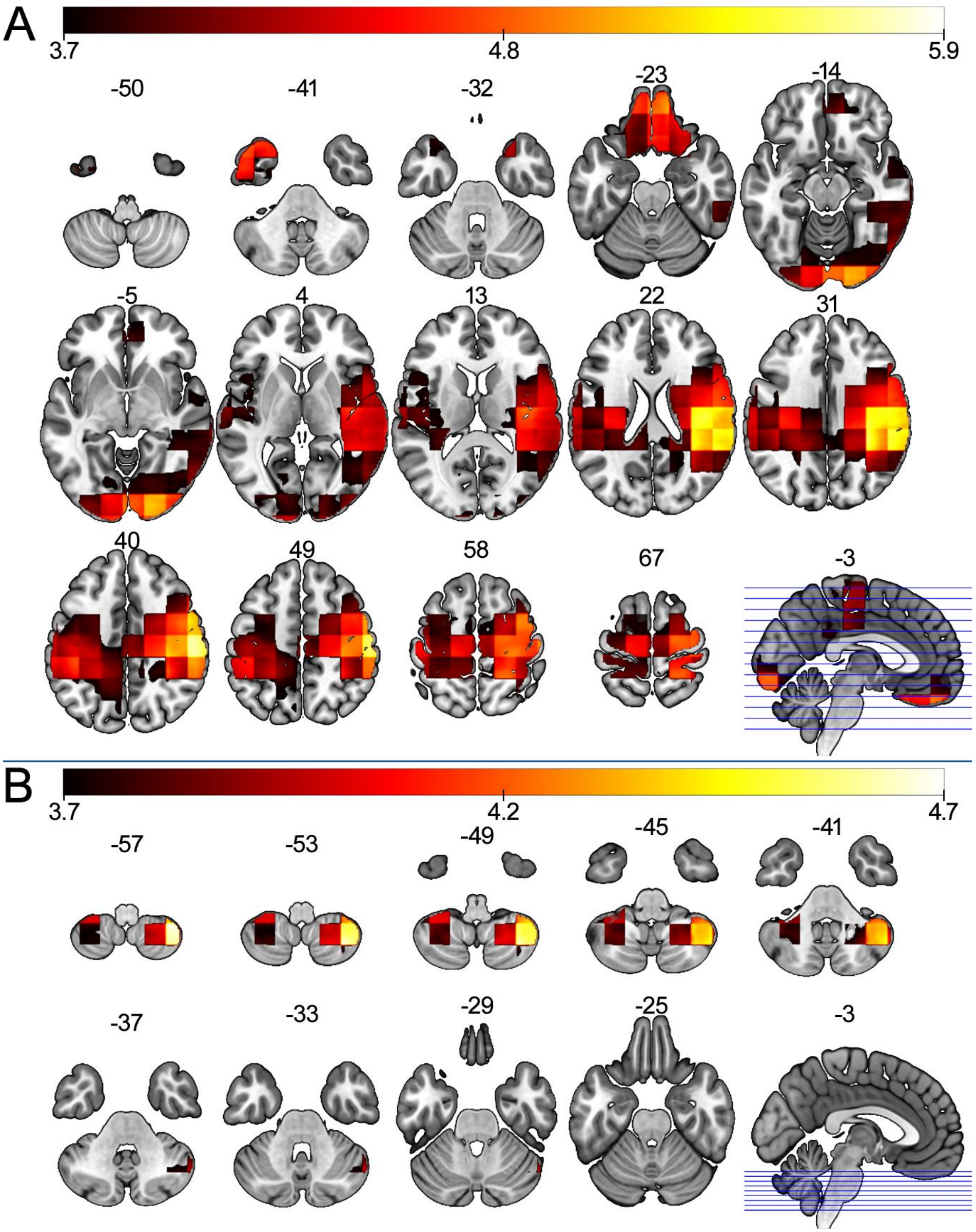

**Supplemental Figure S5. Voxel-wise association between Black race (Panel A), poor sleep quality (Panel B) and L-brainAGE in males.** Results are significant at family-wise error corrected $P_{FWE} < .05$.

#### 6. Associations between G-brainAGE and functional outcomes

**Supplementary Table S6. Spearman rank correlations for functional correlates of G-brainAGE**

| | Females | | Males | |
|---|---|---|---|---|
| | ***rho* (95% CI)** | ***p* (95% CI)** | ***rho* (95% CI)** | ***p* (95% CI)** |
| Endurance | -0.10 (-0.11, -0.10) | 0.10 (0.08, 0.12) | -0.05 (-0.06, -0.04) | 0.43 (0.38, 0.49) |
| Gait speed | -0.02 (-0.03, -0.01) | 0.60 (0.55, 0.65) | -0.05 (-0.06, -0.05) | 0.40 (0.35, 0.44) |
| Grip strength | -0.01 (-0.01, 0.00) | 0.67 (0.63, 0.72) | -0.03 (-0.04, -0.03) | 0.58 (0.53, 0.63) |
| Fluid intelligence | 0.00 (-0.01, 0.00) | 0.67 (0.63, 0.71) | 0.02 (0.01, 0.02) | 0.67 (0.63, 0.71) |
| Crystalized intelligence | -0.06 (-0.06, -0.05) | 0.34 (0.30, 0.39) | -0.05 (-0.06, -0.05) | 0.40 (0.35, 0.44) |

To account for dependence between observations due to relatedness in the HCP data, stratified bootstrapping was carried out with 100 iterations such that each sample consisted of unrelated individuals (i.e., one randomly selected participant per family, n = 445). The reported statistic is the bootstrap distribution mean and 95% confidence intervals (CI).
